# NanoDesigner: Resolving the complex–CDR interdependency with iterative refinement

**DOI:** 10.1101/2025.02.25.640028

**Authors:** Melissa Maria Rios Zertuche, Şenay Kafkas, Dominik Renn, Magnus Rueping, Robert Hoehndorf

## Abstract

Camelid heavy-chain only antibodies consist of two heavy chains and single variable domains (VHHs), which retain antigen-binding functionality even when isolated. The term “nanobody” is now more generally used for describing small, single-domain antibodies. Several antibody generative models have been developed for the sequence and structure co-design of the complementary-determining regions (CDRs) based on the binding interface with a target antigen. However, these models are not tailored for nanobodies and are often constrained by their reliance on experimentally determined antigen–antibody structures, which are labor-intensive to obtain. Here, we introduce NanoDesigner, a tool for nanobody design and optimization based on generative AI methods. NanoDesigner integrates key stages — structure prediction, docking, CDR generation, and side-chain packing — into an iterative framework based on an expectation maximization (EM) algorithm. The algorithm effectively tackles an interdependency challenge where accurate docking presupposes *a priori* knowledge of the CDR conformation, while effective CDR generation relies on accurate docking outputs to guide its design. NanoDesigner approximately doubles the success rate of *de novo* nanobody designs through continuous refinement of docking and CDR generation.

## 1 Introduction

Antibodies are key components of the adaptive immune system, responsible for recognizing and neutralizing antigens. Structurally, they are Y-shaped proteins with two identical heavy and light chains, each containing variable and constant domains. Camelid heavy-chain only antibodies consist of two heavy chains and single variable domains (VHHs) [1], which retain antigen-binding functionality even when isolated. The term “nanobody”, trademarked by the company Ablynx (Ghent, Belgium), is now more generally used for describing small, single-domain antibodies. Due to their high specificity and low immunogenicity, these immunoglobulins are important in research and therapeutic applications [2]. Their characteristics are largely attributed to the unique structural features of the variable domains, composed of four framework regions and three hypervariable complementarity-determining region (CDR) loops. Collectively, these loops form the paratope — the site that interacts with the antigens. While most CDRs adopt constrained canonical conformations, the third CDR loop of the heavy chain (CDRH3) stands out for its high mutability. This variability significantly enhances the diversity of the antibody repertoire, enabling it to bind a wide variety of antigens [3].

Since the regulatory approval of the first monoclonal antibody (mAb) therapy in 1986, over 100 antibody therapies have been introduced globally [4]. In contrast, only four VHH-based drugs have been approved since the first one in 2019 [2]. Nanobodies may offer significant advantages, including enhanced solubility, tissue penetration, and the ability to be produced in a wider range of expression systems, all while effectively targeting similar epitopes as conventional antibodies. Their smaller paratope is compensated by greater structural diversity, especially in the CDRH3 region, which is typically 3 to 4 residues longer than in conventional antibodies [5, 6]. Additionally, an extra disulfide bond between the CDRH1 and CDRH3 regions of the VHHs enhances rigidity and stability compared to conventional antibodies [7]. These structural features enable VHHs to maintain their functionality under harsh conditions, such as high temperatures and denaturant environments, making them effective in situations where traditional antibodies might lose activity, broadening their range of potential applications [7].

Most drug candidates in clinical trials are still developed using traditional *in vitro* techniques, which are time-consuming and resource-intensive [8]. The increasing availability of antibody-specific data has led to the development of bioinformatics tools for rational antibody design [9]. While comprehensive computational workflows for antibody design and optimization have been reported as well, and have even been designed automatically based on interacting agents based on Large Language Models (LLMs) [10], they often depend on computationally demanding methods such as energy minimization and molecular dynamics [11–14]. These approaches may struggle with local optima and constrained search spaces, primarily due to the extensive sampling required. The search space for a CDR of length *L* is vast, with up to 20^*L*^ possible sequences, making exhaustive exploration impractical. To navigate this space efficiently, generative machine learning models have been developed to learn patterns from large datasets, offering improved accuracy and efficiency in predictions while reducing computational overhead [15–18].

Generative models for antibody design initially relied on large language models trained on extensive protein sequence datasets to suggest potential mutations [19, 20]. However, these approaches are limited by their inability to incorporate essential structural information, which is crucial for accurately modeling the 3D nature of proteins. To address this limitation, methods based on Graph Neural Networks (GNNs) integrate both sequence and structural data by representing the entire protein complex (heavy, light, and antigen chains) as a graph [21], effectively capturing spatial and relational dynamics that enhance binding predictions in antibody design. Early efforts to leverage GNNs [17] began by conditioning the CDRH3 design on the antibody framework region, later expanding to include interactions with the epitope [18], albeit only considering the CDRH3 loop and ignoring the rest of the antibody structure. Subsequent advancements introduced equivariant GNNs that account for the rotational and translational symmetries inherent in protein structures, achieving state-of-the-art performance [22]. Similarly, diffusion-based models have advanced the field by enabling the co-design of CDR sequences and their corresponding structures, conditioned on the entire antibody–antigen complex, while also incorporating information about the orientation of the amino acids side chains within the binding interface [15, 23]. Reflecting the growing interest in nanobody design, concurrent work has finetuned RoseTTAFold Diffusion (RFDiffusion) for single-domain antibody generation [24], demonstrating the adaptability of diffusion models to this subclass of antibodies. A recently published review explores recent computational advancements in nanobody research, their impact on antigen-binding and conformational dynamics, and presents a practical example of improving a nanobody-based immunosensor, outlining future directions in healthcare and diagnostics in more detail [25].

Methods trained on antibody–antigen complex structures aim to learn the interactions within binding interfaces, theoretically enabling the *de novo* design of antibodies targeting unseen antigens. However, most generative methods for antibody design, including those adapted for nanobody generation, do not explicitly address the structural interdependencies between docking and CDR design. The generative CDR design process assumes prior knowledge of the antibody–antigen complex, which, when unavailable, is typically derived through computational docking. Yet, generating protein complexes via docking requires knowledge of the antibody’s structure, including the CDR. This interdependency — where CDR design relies on the complex, and complex generation depends on the CDR — remains an unresolved challenge in both antibody and nanobody design.

We developed **NanoDesigner** (see Figure 1), a novel method designed for the *de novo* design as well as the optimization of nanobodies. NanoDesigner integrates key stages in nanobody design, including structure prediction, docking, CDR generation, and side-chain packing. While similar workflows for antibody design are linear, we use a form of expectation maximization (EM) [26] to address the challenge that CDR design depends on accurate complexes and generating complexes requires a CDR. Our algorithm specifically improves the precision of CDRH3 loop predictions and maximizes the likelihood for identifying high-affinity, diverse nanobody candidates, over several baselines. NanoDesigner is freely available at https://github.com/bio-ontology-research-group/NanoDesigner.

**Fig. 1:**
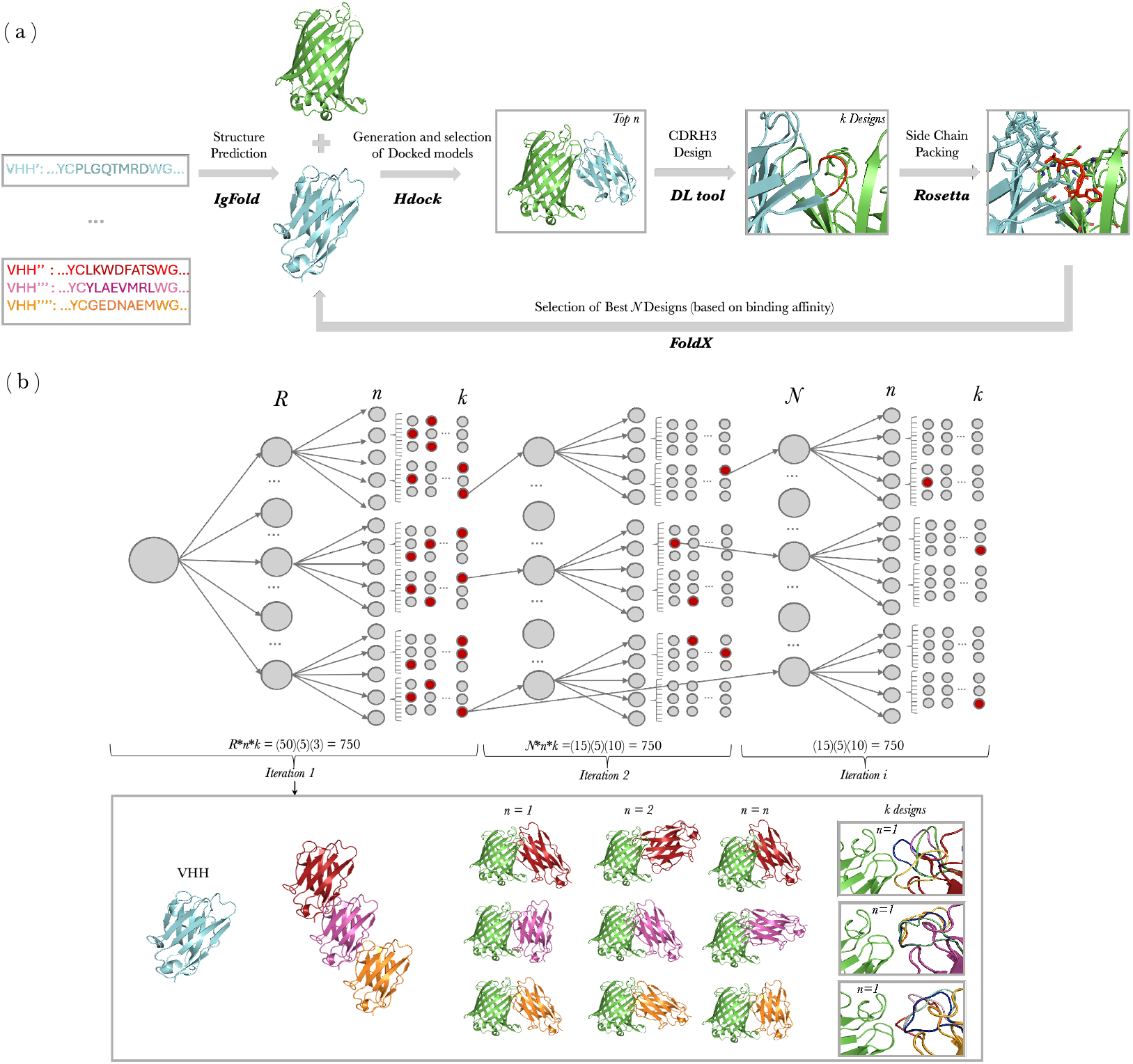
NanoDesigner workflow for the *de novo* design and optimization of nanobodies. Panel (A) shows the workflow; panel (B) displays the variable values and total number of designs across iterations.

## 2 Methods

### 2.1 Datasets

The data used in this study was retrieved from the Structural Antibody Database (SAbDab) in February 2024 [27]. The initial dataset included 5,852 antibody–antigen complexes, of which 1,375 are VHH antibody–antigen complexes. We filtered the entries to retain those with a resolution below 4 Å, targeting protein or peptide antigens, and exhibiting CDRH3–antigen interactions. To further refine the dataset, we ensured that distinct binding interfaces within the same PDB entry were retained, providing diverse interaction information for model training. See Section 2.3 for details on interface determination methods. The expanded dataset was then divided into two sets: one consisting solely of nanobodies, and the other combining nanobodies with light-chain-containing antibodies. For model training, validation, and testing, we clustered the CDRH3 sequences using MMseqs2 [28] with a 40% sequence identity threshold, following prior work [17, 29], with sequence similarity assessed using the BLOSUM62 substitution matrix [28]. This process resulted in 120 clusters for antibodies, 548 for nanobodies, and 668 for the combined dataset. Additionally, to assess the ability of the models to generalize to novel antigens, we employed another clustering approach based on antigen sequence similarity, using a 95% similarity threshold. This method resulted in 177 clusters for antibodies, 804 for nanobodies, and 981 for the combined dataset.

We split the clusters into 10 folds. For 10-fold cross-validation, we trained on eight folds, used one fold for validation, and one for testing, iterating for 10 times and using all 10 folds for testing once; performance results are averaged across the 10 folds. For some experiments, we did not perform cross-validation due to computational limitations and instead used a single 8/1/1 split (based on the clustering results) for training, validation, and testing. Further details of the dataset statistics are shown in Supplementary Table S1 and Supplementary Figure S1.

### 2.2 Structure prediction, docking, and CDR design

For structure prediction, we employed IgFold which is faster than AlphaFold and supports nanobody modeling [30]. For docking, we chose HDOCK [31] for its efficiency in sampling thousands of models rapidly and its ability to integrate binding site information. For CDR design, we evaluated and applied several methods. ADesigner [32] leverages graph neural networks and represents the CDR as ℛ := {(*s*_*i*_, *x*_*i*,*ω*_) | *i* = *l* + 1, …, *l* + *m* }, where each residue is denoted by it type *s*_*i*_, one of the 20 standard amino acids, and its backbone atom coordinates *x*_*i*_, *ω* ∈ ℝ^3^ for the C_*α*_, N, C, O atoms. DiffAb [33], on the other hand, is a diffusion model that allow atomic-resolution antibody design. It achieves this by modeling each residue by its type, coordinates of the C_*α*_ atom and side-chain orientations, denoted as *s*_*i*_, *x*_*i*_ ∈ ℝ^3^, *O*_*i*_ ∈ *SO*(3), respectively. The CDR is initialized arbitrarily, ℛ := {(*s*_*j*_, *x*_*j*_, *O*_*j*_) | *i* = *l* + 1, …, *l* + *m* }, and the model aggregates information from the antigen and the antibody structure, then each amino acid is updated iteratively. In addition to these methods, we considered dyMEAN [16], an end-to-end deep learning tool capable of performing structure prediction, docking, and CDR generation simultaneously. It achieves full-atom design by handling residue-specific representations in a graph, where each residue is a combination of its type *s*_*i*_ and a multi-channel 3D coordinate matrix 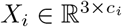, where *c*_*i*_ denotes the number of atoms. dyMEAN incorporates structural initialization and a shadow paratope to guide the docking process, aiming to improve the accuracy of epitope-binding CDR design.

To modify these tools for nanobody design, we made several adjustments. The original input representation in dyMEAN and ADesigner assumed the presence of a light chain. To address this, we adjusted these methods to allow them to handle nanobody structures as well. We make all modifications available on our Github repository at https://github.com/bio-ontology-research-group/NanoDesigner.

As last step, since DiffAb and ADesigner focus only on backbone structures, we perform side-chain packing with Rosetta [34].

### 2.3 Data quality control

For both input and generated data, ensuring the involvement of functional residues in binding, as well as their structural plausibility are critical for accurate antibody and nanobody design. To determine the interacting residues in nanobody- or antibody– antigen complexes, we first extracted the sequence and positional information of the CDRs using the ImMunoGeneTics (IMGT) numbering scheme [35]. This information was then used as input for differential Solvent Accessible Surface Area (dSASA) computations [36], enabling the identification of antigen residues interacting with each of the three CDRs.

To assess structural reliability of the complexes, we quantified both intra- and inter-chain steric clashes, following the approach in [37]. When clashes are detected, we performed refinement through molecular dynamics simulations and energy minimization using the AMBER99 force field within the OpenMM toolkit [38]. This iterative process reduces steric clashes and optimizes molecular geometry by adjusting atomic positions to achieve minimized potential energy.

### 2.4 Evaluation scores

To quantitatively assess the structure and sequence of the generated CDR and the resulting complexes, we use amino acid recovery (AAR), which quantifies the overlap ratio between predicted and actual amino acid sequences, and root mean square deviation (RMSD), which measures structural deviation by comparing the absolute coordinates of the C_*α*_ atoms in the CDR region after Kabsch alignment [39]. Additionally, we used the TM-score [40] and local distance difference test (lDDT) [41] to evaluate global and local structural similarity, respectively, with TM-score focusing on overall topological similarity and lDDT assessing atom-level distance differences. We also use DockQ [42] to evaluate docking quality; DockQ combines scores such as interfacial contact preservation, ligand RMSD, and interface RMSD. DockQ is sensitive to the precise orientation and positioning of the antibody relative to the epitope. Additionally, we calculated differential and relative binding free energy, Δ*G* and ΔΔ*G*, using the FoldX software suite [43], to determine interaction energies and changes in binding affinity due to mutations or perturbations (in *kcal/mol*). Finally, we also report the success rate, defined as the percentage of designs with a negative ΔΔ*G* value. This score quantifies how many designs resulted in improved binding affinity compared to a reference complex. All results are reported as the mean ± a margin of error, corresponding to half the width of the 95% confidence interval, 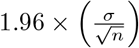.

## 3 Results and Discussion

### 3.1 CDR design for nanobodies

Most existing antibody CDR design tools are trained and tested on light-chain-containing antibodies or mixtures of different types of antibodies, which may limit their applicability to nanobodies. To address this limitation, we developed a nanobody-specific dataset, and an extended dataset that includes both nanobodies and conventional antibodies (see Section 2.1). We clustered both datasets based on antigen sequence similarity, and we also clustered each dataset based on the CDRH3 sequence similarity. We use these clusters to generate training, validation, and test sets; the aim of clustering is to ensure that training, validation, and test sets all contain distinct molecules, i.e., that there is no overlap between training and testing data.

We used 10-fold cross-validation (where folds are generated from the clusters) to evaluate CDR design models in four configurations (based on a dataset containing either only nanobodies or nanobodies plus antibodies, and based on clustering by antigen sequence similarity as well as by CDRH3 sequence similarity). These four strategies allow us to determine whether model training benefits from including light-chain-containing antibodies (i.e., whether there is a transfer of knowledge from antibodies to nanobodies), and which split will generalize better. Table 1 shows the results of our experiments.

**Table 1:**
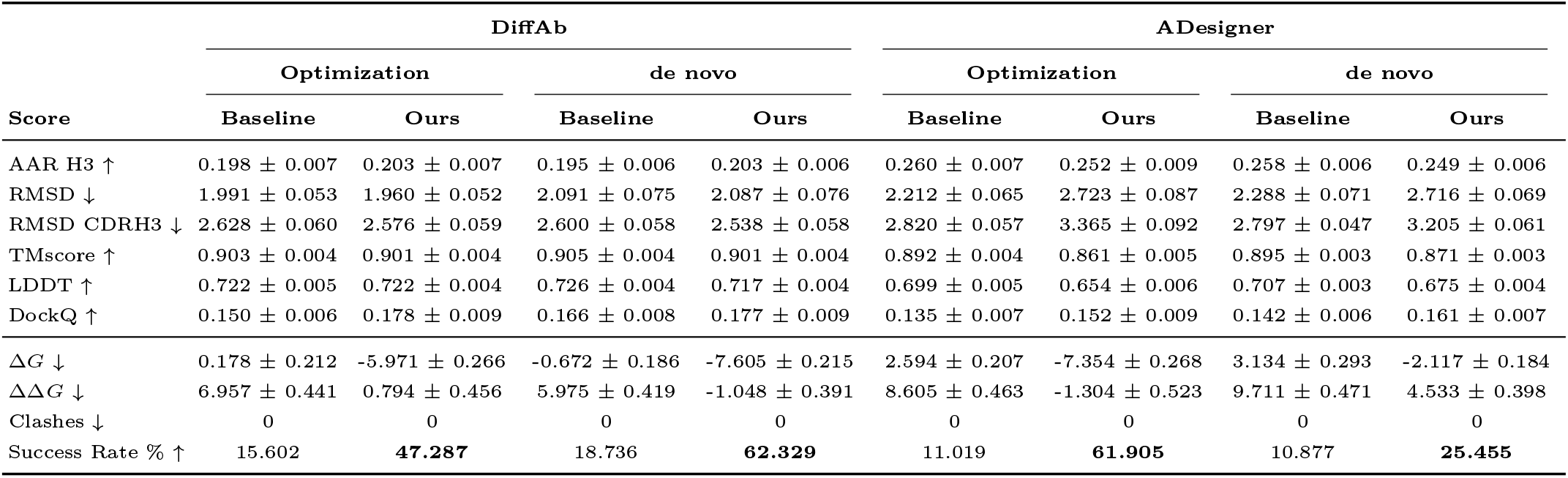
Assessment of CDR antibody design tools on nanobody CDRH3 sequence and structure co-design using 10-fold cross-validation. Performance measures are reported as average ± margin of error obtained from each trained on two datasets: dataset 1 contains only nanobodies, while dataset 2 comprises a mix of nanobodies and antibodies. Both datasets are clustered according to either antigen or CDRH3 sequence similarity (see Section 2.1) . Bold values indicate the best performance for each score, while underlined values represent the second-best performance.

We find that antibody CDR design tools can be adapted for CDRH3 design in nanobodies. The diffusion-based method DiffAb performed better compared to other methods across most scores (see Table 1 and Supplementary Figure S2). Specifically, DiffAb produced stable structures with fewer atomic clashes (i.e., spatially overlapping atoms between the two structures) than other methods, and it generated designs closely matching the natural distribution of affinity values (Supplementary Figure S3). These results indicate that DiffAb can approximate natural nanobody–antigen interactions. ADesigner performed with slightly lower evaluation metrics but overall comparably to DiffAb. The dyMEAN method showed more variability, likely due to a higher number of atomic clashes in its designs. We also find that the inclusion of light-chain-containing antibodies in the training data did not significantly improve model performance. Finally, clustering and splitting training and testing data by the antigen sequence similarity rather than clustering and splitting by CDRH3 similarity generally yielded better results.

While our results demonstrate that CDR design methods intended for antibodies can design nanobody CDRH3 regions with a performance comparable to those reported in the antibody literature, additional challenges arise when integrating them into a complete design workflow. In particular, the input to the CDR design methods is a complex consisting of the antigen and the nanobody (or antibody). These complexes can be obtained through experimental methods or they can be generated computationally. Computational generation requires knowledge or prediction of the structure of the antigen and the nanobody (including its CDRH3 region). However, design of a CDRH3 will depend on the complex in the sense that, if the complex is different (e.g., if the nanobody binds at a different epitope), the designed CDRH3 will also be different. This leads to a mutual dependency: the design of the CDRH3 region depends on the complex, and computational generation of the complex depends on the nanobody structure (in particular its CDRH3 region at which it binds).

### 3.2 NanoDesigner: End-to-end design and optimization

We developed **NanoDesigner**, an algorithm that addresses the mutual dependency between protein complex generation and CDRH3 design using Expectation-Maximization (EM) [44]. The algorithm consists of iterating between two steps: in the E-step, we generate a candidate protein complex given the current best guess of a CDRH3 region; in the M-step, we design or optimize a CDRH3 region given the generated complex. We iterate between the E- and M-steps multiple times, and further combine this EM method with a beam search where we keep the top *n* ranking designs after each M-step. The NanoDesigner algorithm is described in detail in the Supplementary Materials.

NanoDesigner takes as inputs the structures of a nanobody as scaffold, an antigen, and the epitope information. During the initialization step, the VHH sequence is extracted, and since the CDRH3 is unknown at this stage, we randomize it *r* times to generate *r* VHH sequences with a random CDRH3. NanoDesigner then predicts the structure of each perturbed sequence using IgFold [30].

In the E-step of the algorithm, each of the *R* predicted structures is docked with the antigen structure using HDOCK [31] to generate protein complexes. The location of the epitope and the paratope are provided as additional information to guide the docking process. For each of the randomized nanobodies, HDOCK generates *d* complexes. NanoDesigner then assesses each of these complexes for atomic clashes [37] and determines interacting residues [36].

We remove all complexes that contain clashes after refinement, and models where the CDRH3 or the epitope residues are not involved in the interactions (see Section 2.3). NanoDesigner then ranks all remaining complexes based on epitope recall, using the measure 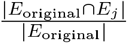 (where *E*_original_ represents the desired epitope, i.e., the input to NanoDesigner, and *E*_*j*_ represents the predicted epitope in the generated complex). The top *n* docked models are then selected to move forward in the process.

In the M-step of the algorithm, a generative model (either DiffAb or ADesigner) is employed to generate *k* designs with novel CDRH3 loops for each nanobody complex. After this step, NanoDesigner has generated a total of *r* × *n* × *k* nanobodies, each in complex with an antigen, and each with a designed and optimized CDRH3. These mutants then undergo side-chain packing using the Rosetta software [34]. The designed nanobodies are then ranked based on the predicted binding affinity energy *δ* computed with FoldX [43], with stronger binding affinities receiving higher rankings. We retain the top-*n* ranking nanobodies and use them as input to the next E-step. Only in the first iteration, we select the top-*n* from all *r* lineages, i.e., we keep the top-ranking nanobody from each of the original *r* (randomized) lineages; in subsequent iterations, we rank all designed nanobodies by binding affinity and keep the top-*n* across all. The reason for keeping the best performing nanobody from each lineage in the first iteration is that we found that binding affinities generally improve more in the second iteration, and keeping each lineage allows us to maintain higher diversity. Figure 1 gives an overview of NanoDesigner.

There are two applications for nanobody CDRH3 design: optimizing the CDRH3 of an existing nanobody with known binding to an antigen (“nanobody optimization”) and designing a CDRH3 *de novo* without any prior information about a complex. NanoDesigner can be used both for nanobody optimization and *de novo* design, by modifying only the final ranking based on the measure *δ*. In nanobody optimization, where a complex is already known, the algorithm optimizes the predicted relative binding free energy (ΔΔ*G*), with the objective to reduce this energy when compared with the reference complex. This approach is useful for enhancing the binding affinity of an existing nanobody, such as one targeting a mutating antigen. In *de novo* design, where no complex is known, the focus is on optimizing the predicted absolute binding free energy (Δ*G*) which is valuable for targeting novel antigens. We compared NanoDesigner with a linear design workflow without the iterative EM loop as a baseline; results are shown in Table 2.

**Table 2:**
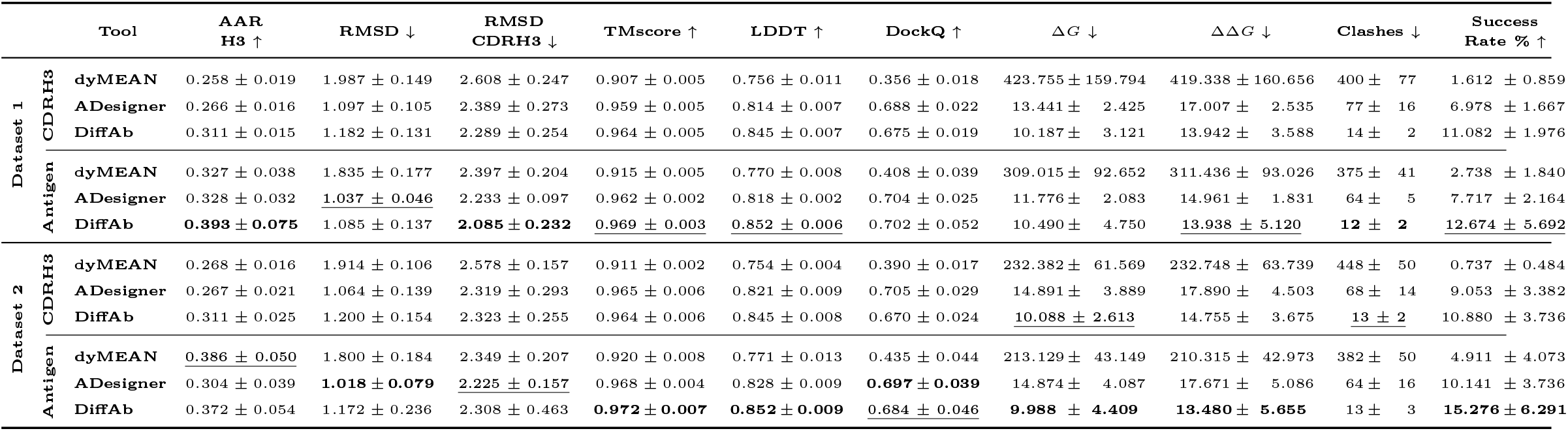
NanoDesigner in the nanobody optimization and *de novo* scenarios. Each pair of columns compares the average ± margin of error for measures obtained on a test set. We compare a linear workflow of NanoDesigner without the EM loop (baseline) with the results of NanoDesigner with the EM loop for 10 iterations. The test set is a single fold from Dataset 1 (consisting of only nanobodies), clustered by antigen sequence similarity (see Section 2.1).

Our results demonstrate that NanoDesigner consistently and substantially improves over the linear design workflow, demonstrating that our iterative refinement approach is successful: NanoDesigner achieves a significantly higher success rate, more than twice compared to the baseline linear workflow, and achieves, for example, 62% success rate using DiffAb and 25% using ADesigner for CDRH3 design. This improvement is driven by iterative optimization, which refines nanobody designs and enhances binding affinities, as shown in Supplementary Material Figure S4, where ΔΔ*G* values steadily improve across iterations. Although NanoDesigner shows slightly lower performance on CDRH3 design scores such as AAR, TMScore, LDDT, and DockQ (Table 1), it substantially improves binding energy. The improved Δ*G* and ΔΔ*G* values suggest that greater diversity in the CDRH3 region drives stronger affinities. By exploring a wider range of conformations, NanoDesigner enhances binding and reduces the likelihood of clashes, leading to more effective interactions with the target antigen. Similar trends were observed in the *de novo* design scenario, with NanoDesigner showing consistent improvements in binding affinity compared to the baseline. As shown in Table 2, our method enhances success rates and binding energetics, demonstrating its robustness across both optimization and *de novo* design tasks.

In Supplementary Material Table S2, we present the performance of dyMEAN CDRH3 design on a reference test set, as well as on the same *r* randomized complexes used by NanoDesigner in both the DiffAb and ADesigner evaluations. All scores remained similar across the evaluated datasets. In terms of energy scores and structural clashes, dyMEAN has a high variability and substantially lower success rates. This may be due to some limited overfitting and a higher frequency of structural clashes, ultimately limiting its effectiveness as a nanobody design tool. Our findings are consistent with a recent study [45], which concluded that dyMEAN has limitations when designing antibodies. We also evaluated the diversity of sequences of the CDR design methods on their own (Supplementary Material Figure S6) and find that dyMEAN generates CDRs with lower diversity than ADesigner and Diffab.

### 3.3 NanoDesigner: Test cases

Based on our findings, we employed NanoDesigner, with DiffAb for the CDRH3 generation stage, on three distinct antigens: mNeonGreen, KRAS, and HER2, selected for their significance in research and therapeutic applications. For mNeonGreen (`PDB:5LRT`) and KRAS (`PDB:4OBE`), epitope sequences were provided by domain experts. For HER2, we identified epitope sequences via dSASA analysis using HER2’s antibody-bound structure (`PDB:8PWH`); in this structure, trastuzumab and pertuzumab define two distinct epitopes. The HER2 antigen chain was then isolated for use in the NanoDesigner process (see Section 2.3).

Our analysis of the complexes generated by NanoDesigner on our evaluation datasets showed that, usually, all three CDRs are highly involved in binding (see Supplementary Material Figure S1, panel (b)). Therefore, for the three test cases, we designed all three CDRs instead of only CDRH3; we trained DiffAb for CDR design, and used NanoDesiger for the *de novo* prediction and optimization scenarios (see Supplementary Table S3 and Supplementary Figure S5).

We used NanoDesigner to design either one (CDRH3) or all three CDRs for each of the three targets. Figure 2 indicates that, when focusing only on CDRH3, the algorithm converges more quickly and with better Δ*G* values, and with a steeper decline in energy during early iterations compared to models that optimize all three CDRs. Additionally, the variability of Δ*G* values across our test cases suggests that some epitopes, such as the epitopes we selected in KRAS and HER2, present greater difficulty in design, likely due to more complex or variable binding interfaces.

**Fig. 2:**
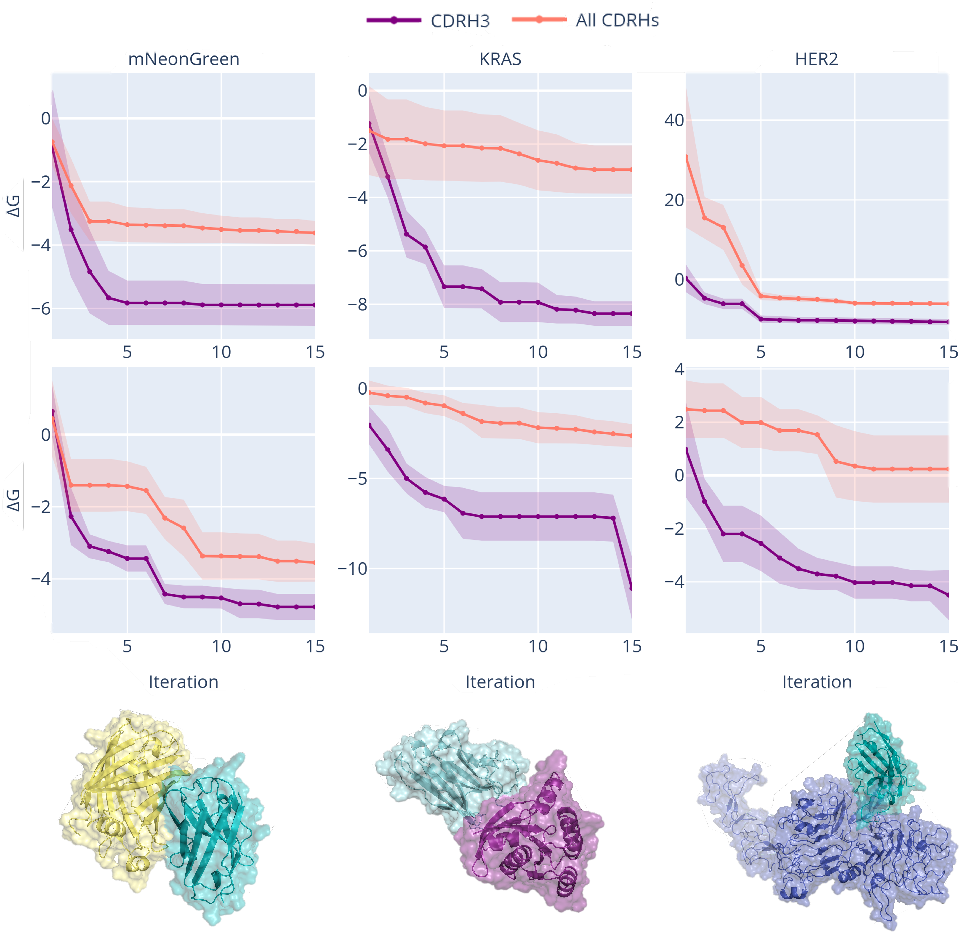
NanoDesigner for *de novo* design — test cases. The Δ*G* values for the design of CDRH3 (purple) or all CDRs (orange) are tracked across iterations for three distinct antigens: mNeonGreen (`PDB:5LRT`), KRAS (`PDB:4OBE`), and HER2 (`PDB:8PWH`), represented from left to right in the panels. A camelid nanobody scaffold (`PDB:6LR7`) was utilized in the first two panels, while a humanized nanobody scaffold (`PDB:7EOW`) was employed in the third panel. Within each antigen panel, two distinct epitope sites were analyzed. The first epitope is displayed at the top, while the second is shown in the middle. At the bottom of each panel, a representative design per nanobody– antigen complex is illustrated for the first epitope. The plotted lines represent average Δ*G* values across epitopes, with shaded regions indicating variability. Epitope details and VHH scaffold data are available in Supplementary Table S4.

## 4 Conclusion

We developed NanoDesigner, an algorithm that improves the *de novo* nanobody design as well as nanobody optimization. NanoDesigner overcomes a limitation in traditional antibody design methods where generation of CDR loops using generative models is dependent on knowledge of a complex, and predicting the complex structure depends on knowing the CDR sequence. NanoDesigner also provides specific optimizations and evaluation results for nanobodies, in contrast to other methods focusing on general antibodies. NanoDesigner is suited for designing nanobodies for previously uncharacterized antigens and antigens for which neither a nanobody nor antibody is available.

## Supplementary information

This article is accompanied by a Supplementary Materials file that provide detailed additional analyses, extended data tables, and supplementary figures that further support the findings discussed in the main text.

## Declarations

### Ethics approval and consent to participate

Not applicable.

### Materials Availability

Not applicable.

### Code Availability

The code supporting the findings of this study is freely available on https://github.com/bio-ontology-research-group/NanoDesigner. All data generated or analyzed in this study can be reproduced by following the procedures detailed in the link. Users can access the code and replicate the steps provided to produce the data associated with this study.

### Data Availability

The datasets utilized in this study are available from SAbDab database, accessible at https://opig.stats.ox.ac.uk/webapps/sabdab-sabpred/sabdab. Detailed instructions on accessing and processing these datasets are provided in the our Github repository. An analysis of the employed dataset can be found in the Supplementary Material as well as the ‘Datasets’ section of the manuscript.

### Author contribution

R.H. and M.R. conceived of the study. Ş.K., R.H., M.R., M.M.R.Z. designed the study. M.M.R.Z. implemented the study and performed all primary analysis. D.R. and M.R. contributed chemical expertise to design, implementation, and contributed to analysis. Ş.K., R.H., M.R. supervised the research. R.H., Ş.K., M.R. acquired funding for the study. M.M.R.Z. wrote the initial manuscript. All authors reviewed and revised the manuscript.

## Acknowledgments

We acknowledge support from the KAUST Supercomputing Laboratory. We would like to thank Jörg Eppinger for his thorough contributions and insightful discussions, which greatly aided the development of the workflow.

