## Supplementary Materials for "NanoDesigner: Resolving the complex–CDR interdependency with iterative refinement"

#### Supplementary Material

Melissa Maria Rios Zertuche[1], Şenay Kafkas[2], Dominik Renn[3], Magnus Rueping[3,4,5,6], Robert Hoehndorf[1,4,5,7,8,*]

[1]Biological and Environmental Science and Engineering (BESE) Division, King Abdullah University of Science and Technology, 23955-6900, Thuwal, Saudi Arabia.

[2]KAUST Academy, King Abdullah University of Science and Technology, 23955-6900, Thuwal, Saudi Arabia.

[3]KAUST Catalysis Center (KCC), Division of Physical Sciences and Engineering, King Abdullah University of Science and Technology, 23955-6900, Thuwal, Saudi Arabia.

[4]KAUST Center of Excellence for Smart Health (KCSH), King Abdullah University of Science and Technology, 23955-6900, Thuwal, Saudi Arabia.

[5]KAUST Center of Excellence for Generative AI, King Abdullah University of Science and Technology, 23955-6900, Thuwal, Saudi Arabia.

[6]Institute for Experimental Molecular Imaging (ExMI), University Clinic, RWTH Aachen, Forckenbeckstraße 55, D-52074, Aachen, Germany.

[7]SDAIA–KAUST Center of Excellence in Data Science and Artificial Intelligence, King Abdullah University of Science and Technology, 4700 King Abdullah University of Science and Technology, Thuwal, Saudi Arabia.

[8]Computer, Electrical and Mathematical Sciences and Engineering Division, King Abdullah University of Science and Technology, 23955-6900, Thuwal, Saudi Arabia.

### S1 Dataset statistics and analysis

The data used in this study was retrieved from the Structural Antibody Database (SAbDab) in February 2024 [1]. The initial dataset included 5,852 antibody–antigen complexes, of which 1,375 are VHH antibody–antigen complexes. Instances in the dataset correspond to entries in the Protein Data Bank (PDB) [2]. The PDB entries contain structural information for all chains present, including the identifiers for heavy, light, and antigen chains, and relevant metadata such as the resolution of the structure. Instances were filtered to retain only those with a resolution quality below 4 Å, retain nanobodies that target protein or peptide type antigens, and ensure that the heavy, light, and antigen chain identifiers were completely identified. Given that the generative models used in our work are designed to learn from interactions involving the complementarity-determining region (CDR) loops of the heavy chain, we removed all complexes that did not have interactions with the CDR loops; we calculated the delta solvent accessible surface area (dSASA) to determine the binding interface [3]. All entries were renumbered according to the IMGT scheme, which assigns specific positions based on the conserved residues in the antibody framework. This provided a standardized way to locate the CDRs, with the paratope defined as the residues comprising the CDRH1, CDRH2, and CDRH3 loops. We used dSASA calculation to generate an interaction matrix between two biomolecules and identify specific binding pairs, and we used this to compute the percentage involvement of each CDR. All filtered entries showing interaction were renumbered back to their appropriate numbering schemes to ensure compatibility with the different generative tools used in subsequent analyses. For instance, Chothia numbering [4] was applied for DiffAb. As seen in Table S1, despite having 1,375 unique PDB identifiers for nanobodies, we identified 1,678 unique binding interfaces after data pre-processing. The reason for this increase is that some PDB complexes have multiple nanobodies binding to the antigen, and using different epitopes. In the case of light-chain-containing antibodies, while there are 4,477 unique PDB complexes, the number of unique binding interfaces is substantially lower, with only 283 instances involving interactions from CDRH3. This is likely due to different binding mechanisms involving the light chains or other effector regions. This difference highlights the distinct binding behaviors and structural constraints between light-chain-containing antibodies and nanobodies in targeting antigens, which is a crucial consideration for model training and evaluation in our study.

**Table S1**: Dataset statistics. The number of unique PDB complexes is shown in parenthesis; total number of instances refers to different nanobody– or antibody–antigen pairs.

| | **Antibodies** | **Nanobodies** |
|---|---|---|
| **Total PDB entries** | 4477 (4477) | 1375 (1375) |
| **Total Instances** | 8693 (4477) | 2413 (1375) |
| **Filtered instances** | 3113 (1396) | 1793 (945) |
| **Instances interacting from the paratope** | 285 (171) | 1709 (933) |
| **Instances interacting from CDRH3** | 283 (169) | 1678 (921) |

All nanobody instances retained after the data pre-processing step were analyzed to determine the lengths of CDRH1, CDRH2, and CDRH3, as well as their average involvement in interactions with corresponding antigen proteins or peptides.

Panel (a) from Figure S1 displays the length distributions of CDRH1, CDRH2, and CDRH3. The lengths of CDRH1 and CDRH2 show a clear clustering around 8

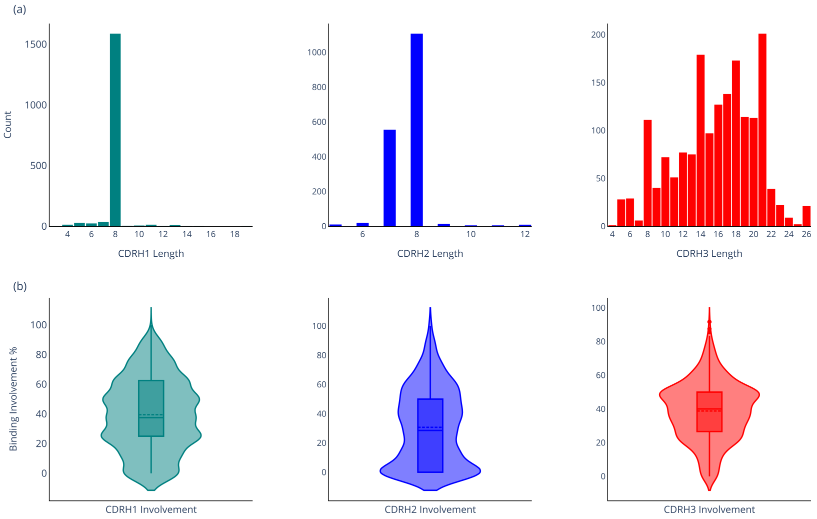


**Fig. S1**: Length and binding involvement of nanobody CDRHs.

amino acids, which may be associated with their tendency to adopt canonical structures commonly observed in nanobodies. In contrast, CDRH3 exhibits a much wider range of lengths, from 4 to over 26 amino acids, indicating substantial variability. This diversity is expected, as CDRH3 plays a critical role in antigen specificity and frequently deviates from canonical shapes, allowing for greater flexibility and adaptability in binding interactions. Panel (b) in Figure S1 shows the binding involvement percentages of CDRH1, CDRH2, and CDRH3 in nanobody–antigen complexes. The violin plots highlight the variability in contribution of each CDR loop to antigen binding. CDRH1 and CDRH2 exhibit moderate involvement, with median values around 40-50%, indicating their consistent, but less dominant, roles in binding interactions. In contrast, CDRH3 shows a broader range of involvement, with a high frequency of instances near the upper bound of participation, supporting its role as the primary driver of antigen specificity. The variability in CDRH3 binding involvement aligns with its structural flexibility and longer length, emphasizing its adaptability and key function in facilitating diverse immune responses.

### S2 Assessment of generative methods for nanobody CDRH3 design

#### S2.1 CDRH3 design for nanobodies

We evaluated and benchmarked each CDR design method individually, outside the NanoDesigner workflow. Figure S2 visually presents the distribution of key metrics, including amino acid recovery for CDR3 (AAR H3) and structural performance indicators such as root mean square distance (RMSD) and TM-score, comparing the generated designs to a reference nanobody-antigen complex. DiffAb consistently outperformed the other methods across most metrics, demonstrating lower variability and better structural alignment with the reference. ADesigner followed closely, although with slightly higher variability, while dyMEAN exhibited the greatest variability, indicating inconsistent CDRH3 reconstruction. The RMSD metric for the CDRH3 loop (third panel, top row) after Kabsch alignment [5] deviates from this trend, and DiffAb does not significantly outperform the other methods in this case.

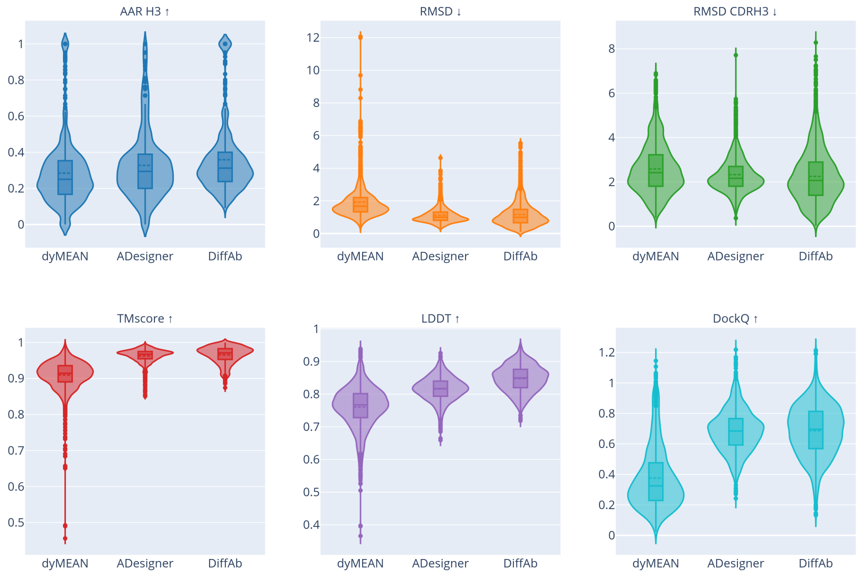


**Fig. S2**: Performance comparison of CDR design tools in nanobody CDRH3 prediction.

#### S2.2 Affinity distribution of generated nanobodies

Figure S3 shows a comparison of the differential binding free energy ($\Delta G$, in $kcal/mol$) distributions between the original nanobody dataset (second column, last row of Table S1) and the pooled designs generated by DiffAb, ADesigner, and dyMEAN, all evaluated using the FoldX software suite [6]. The density plots show that DiffAb and ADesigner produce designs that closely match the natural distribution of affinity energies in the dataset, whereas dyMEAN displays a more variable distribution that does not align well with the affinity profiles of the reference dataset.

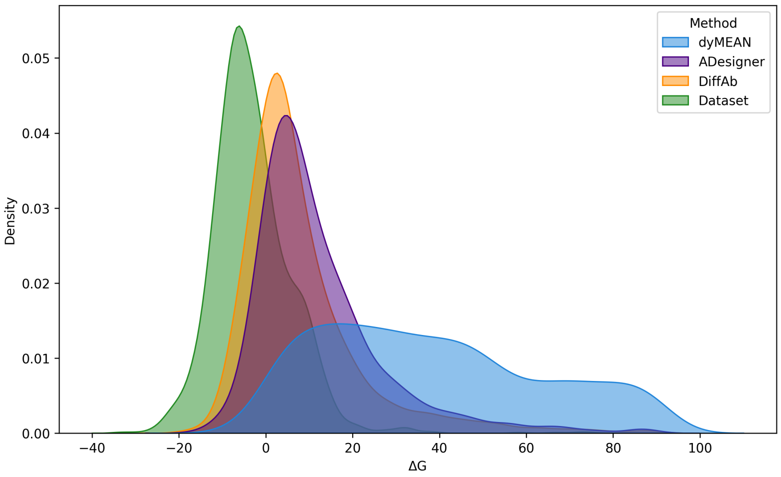


**Fig. S3**: Affinity distribution of natural and predicted nanobody-antigen complexes across CDR design tools.

### S3 NanoDesigner algorithm description

The NanoDesigner algorithm is divided into key phases analogous to the steps in the expectation–maximization (EM) algorithm: initialization, expectation, maximization, and convergence test. In the initialization phase, the nanobody scaffold undergoes sequence randomization, with $r = 50$ sequences being generated in our experiments. This randomization focuses on either the CDRH3 or all CDRs simultaneously, serving as an initial exploration of the sequence space to generate candidates for subsequent design and optimization. For structure prediction, IgFold is used to predict 3D structures for the $r$ sequences. Throughout the workflow, quality control checks, such as structural validation and refinement, are integrated at every stage to ensure only reliable intermediate structures proceed to subsequent steps.

During the expectation step, a set of nanobody–antigen complexes is obtained through docking simulations performed by HDOCK for each randomized nanobody sequence. The inputs include the structure of the randomized nanobody, the target antigen structure, and their respective complementary binding site information: the paratope and the epitope. The paratope is defined as the collection of the three CDRs. If a nanobody–antigen complex is provided, the epitope is identified through dSASA analysis during a preprocessing step. Otherwise, the epitope must be provided as an amino acid sequence and mapped onto the antigen structure. HDOCK with default parameters generates 100 candidate models per simulation; this number can be adjusted according to experimental requirements. The complex selection process determines which docked models proceed to the next stage as inputs. Priority is given to complexes that exhibit high epitope recall. To maintain a balance between exploration and optimization, the algorithm employs a beam search strategy, retaining only the top $n$ models sorted by epitope recall. This allows the algorithm to explore a diverse range of solutions while focusing on promising candidates for further refinement and optimization.

In the maximization step, for each docked model, a set of $k$ CDR designs are generated using either the DiffAb or ADesigner methods. These designs undergo a complex selection process to filter out models based on structural quality (presence of "clashes" where atoms would overlap in space) and the involvement of the CDR loops in interactions (i.e., we remove all CDR designs where the generated CDR loop is not involved in an interaction with the antigen epitope). Only the designs that pass these quality filters proceed to the evaluation and ranking phase. The ranking is based on an objective function $\delta$ that measures binding efficiency. The function $\delta$ is either the binding free energy ($\Delta G$) or the relative binding free energy ($\Delta\Delta G$), depending on whether the workflow focuses on *de novo* design or optimization scenarios. At the end of this step, the top $n$ designs with the best ranking scores (based on $\delta$ are selected to proceed to the next iteration.

In all our experiments, convergence is determined by a predefined number of total iterations. This fixed iteration limit ensures that the design process terminates consistently across runs. However, alternative convergence criteria could be applied to halt the iterative process based on change in the overall evaluation metrics.

### S4 Evaluation of nanobody *de novo* design and optimization

#### S4.1 Differential and relative binding free energy trends

NanoDesigner was applied in two distinct scenarios, using two different CDR generation methods (DiffAb and ADesigner), with analysis conducted on one fold. The plots in Figure S4, illustrate the comparison of average predicted $\Delta G$ (free energy change) and $\Delta\Delta G$ (change in binding energy) values over ten iterations across different NanoDesigner configurations. The left panels correspond to the optimization scenario, where the objective function of the algorithm focuses on improving binding affinity compared to a reference complex. The right panels represent the *de novo* scenario, where the algorithm optimizes the absolute binding energy without reference to a pre-existing complex.

In both scenarios, a general trend toward more negative values is observed over the ten iterations, indicating improved binding affinity and energy stability as the process progresses. The boxes show the distribution of the data, with whiskers highlighting the variability across iterations. ADesigner (purple) performs better in the optimization scenario, achieving more negative values consistently, while DiffAb (orange) shows stronger performance in the *de novo* scenario, suggesting it is more effective for generating novel designs without relying on a reference complex.

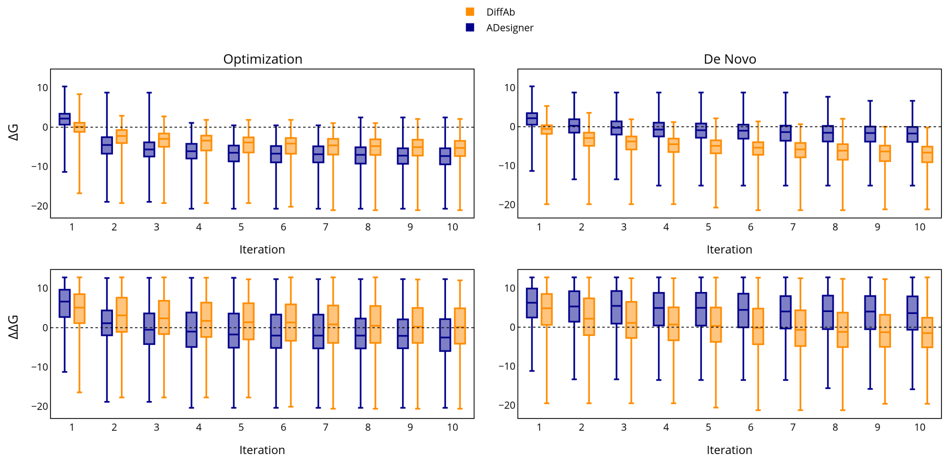


**Fig. S4**: Differential and relative binding free energy across iterations, results from NanoDesigner in the *de novo* and optimization scenarios using two CDR generation methods. Analysis was conducted on one fold from the optimal training configuration using Dataset 1 (consisting of only nanobodies), clustered by antigen sequence similarity.

#### S4.2 Assessment of dyMEAN end-to-end deep learning tool on the *de novo* design and optimization of nanobodies

dyMEAN is an end-to-end model specifically designed for antibody CDR design. Consistent with DiffAb and ADesigner methods, we trained dyMEAN on a nanobody-only dataset, where entries were clustered based on antigen sequence identity. We evaluated dyMEAN's performance on the CDRH3 design task using the same test set of nanobody–antigen complexes. Results were compared against the same complexes with randomized structures and sequences generated during the initialization step of the NanoDesigner workflow (see Section S4). Results are shown in Table S2.

**Table S2**: Performance comparison of dyMEAN across various metrics. Analysis was conducted on one fold using Dataset 1 (consisting of nanobodies only), clustered by antigen sequence similarity. Results are produced based on the same complexes with randomized structures and sequences generated during the initialization step of the NanoDesigner workflow (see Section S3).

| **Metric** | **CDRH3 design** | **with Randomization** |
|---|---|---|
| AAR H3 ↑ | 0.277 ± 0.019 | 0.281 ± 0.020 |
| RMSD ↓ | 2.132 ± 0.288 | 2.249 ± 0.315 |
| RMSD CDRH3 ↓ | 2.824 ± 0.186 | 2.864 ± 0.200 |
| TMscore ↑ | 0.907 ± 0.007 | 0.897 ± 0.013 |
| LDDT ↑ | 0.759 ± 0.010 | 0.749 ± 0.016 |
| DockQ ↑ | 0.390 ± 0.031 | 0.257 ± 0.024 |
| $\Delta G$ ↓ | 160.059 ± 55.907 | 197.617 ± 43.343 |
| $\Delta\Delta G$ ↓ | 166.611 ± 56.117 | 204.093 ± 43.533 |
| Clashes ↓ | 48 ± 28 | 70 ± 22 |
| Success Rate% ↑ | 0.943 | 1.110 |

### S5 Simultaneous design and optimization of all three nanobody CDR loops with NanoDesigner

Based on the performance of DiffAb seen in the CDRH3 design task and when embedded in the NanoDesigner workflow, we conducted further analysis using DiffAb for CDR design. We expanded the evaluation to examine the model's ability to design all three nanobody CDR loops simultaneously. This analysis utilized the same model for both *de novo* design and optimization scenarios within the NanoDesigner framework, employing the same test set as before (Section 3.2).

Table S3 presents the average results and 95% confidence interval $1.96 \times \left(\frac{\sigma}{\sqrt{n}}\right)$. Our findings indicate that CDRH1 and CDRH2 achieved higher average AAR compared to CDRH3, which may be attributed to their shorter lengths and the prevalence of more canonical structures or conserved sequences. Structural measures followed a similar pattern to those observed in the single CDR design experiments. When analyzing energy-related metrics such as $\Delta G$, $\Delta\Delta G$, and the overall success rate, we observed larger improvements with the NanoDesigner iterative loop. Figure S5 further illustrates the trends in energy across iterations, showcasing the iterative refinement achieved through the NanoDesigner framework.

**Table S3**: Assessment of DiffAb in the simultaneous co-design of all nanobody CDRHs alone or within the NanoDesigner workflow for the nanobody *de novo* or optmization scenarios. Analysis was conducted on a unique fold from the optimal training configuration using Dataset 1 (nanobodies), clustered by antigen sequence similarity (See section 2.1). The "baseline" experiments are the NanoDesigner workflow without the iterative refinement loop.

| | | **NanoDesigner on 3CDRHs** | | | |
|---|---|---|---|---|---|
| | **3CDRHs Design** | **Optimization** | | **De novo** | |
| **Score** | **DiffAb** | **Baseline** | **Ours** | **Baseline** | **Ours** |
| AAR H3 ↑ | 0.298 ± 0.006 | 0.160 ± 0.008 | 0.147 ± 0.008 | 0.159 ± 0.009 | 0.127 ± 0.009 |
| AAR H2 ↑ | 0.426 ± 0.009 | 0.176 ± 0.038 | 0.156 ± 0.010 | 0.180 ± 0.012 | 0.178 ± 0.012 |
| AAR H1 ↑ | 0.397 ± 0.010 | 0.256 ± 0.014 | 0.239 ± 0.014 | 0.253 ± 0.015 | 0.128 ± 0.017 |
| RMSD ↓ | 2.508 ± 0.181 | 3.029 ± 0.087 | 3.462 ± 0.082 | 3.115 ± 0.097 | 3.794 ± 0.117 |
| RMSD CDRH3 ↓ | 2.972 ± 0.238 | 3.384 ± 0.079 | 3.487 ± 0.072 | 3.421 ± 0.083 | 3.487 ± 0.089 |
| RMSD CDRH2 ↓ | 1.729 ± 0.035 | 1.707 ± 0.038 | 2.009 ± 0.049 | 1.739 ± 0.040 | 2.615 ± 0.051 |
| RMSD CDRH1 ↓ | 2.546 ± 0.573 | 2.392 ± 0.062 | 2.496 ± 0.063 | 2.426 ± 0.062 | 3.446 ± 0.081 |
| TMscore ↑ | 0.901 ± 0.001 | 0.866 ± 0.002 | 0.844 ± 0.002 | 0.863 ± 0.002 | 0.840 ± 0.003 |
| LDDT ↑ | 0.751 ± 0.002 | 0.660 ± 0.003 | 0.639 ± 0.003 | 0.658 ± 0.003 | 0.640 ± 0.004 |
| DockQ ↑ | 0.514 ± 0.010 | 0.127±0.008 | 0.121±0.008 | 0.127 ± 0.008 | 0.136 ± 0.009 |
| $\Delta G$ ↓ | 25.747 ± 2.062 | 2.221 ± 0.894 | 0.572 ± 0.795 | -0.127 ± 0.173 | -4.041 ± 0.334 |
| $\Delta\Delta G$ ↓ | 32.270 ± 2.115 | 9.331 ± 1.069 | 7.664 ± 1.021 | 7.048 ± 0.710 | 3.802 ± 0.828 |
| Clashes % ↑ | 7 ± 2 | 0 | 0 | 0 | 0 |
| Success Rate % ↑ | 2.747 | 12.593 | 23.810 | 14.375 | 28.736 |

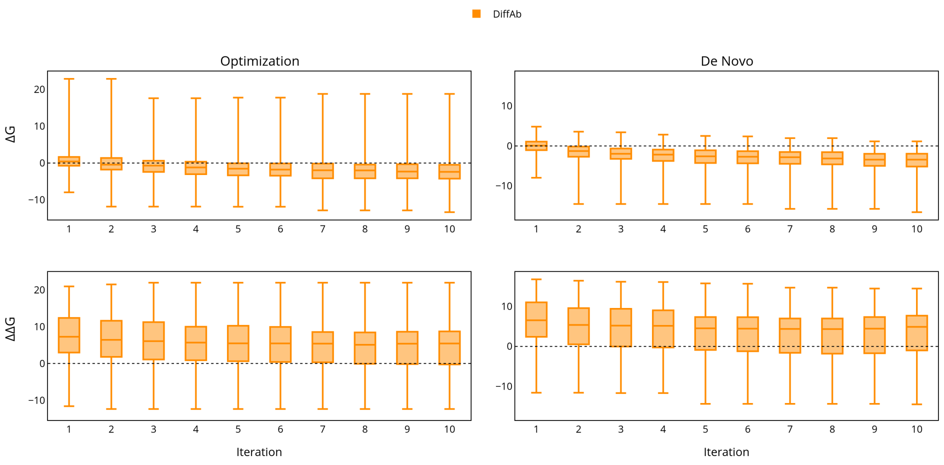


**Fig. S5**: Differential and relative binding free energy across iterations based on NanoDesigner when simultaneously designing all three CDR loops. The evaluation was done on one fold using Dataset 1 (consisting of only nanobodies), clustered by antigen sequence similarity.

### S6 Application of NanoDesigner on specific test cases

We applied NanoDesigner on three specific test cases; the test cases and the epitopes used are shown in Table S4. The epitopes together with the positional information of the three CDR loops were used as input.

**Table S4**: Inputs used for three NanoDesigner test cases. Epitope positional information for mNeon-Green (`PDB:5LRT`) and KRAS (`PDB:4OBE`) was determined using sequence data provided by domain experts. For HER2 (`PDB:8PWH`), epitope residue positions were obtained from the antibody-bound state using dSASA analysis.

| **Source PDB** | | **User Epitope Sequence Input** | **Epitope Extracted Positional Information** |
|---|---|---|---|
| **Nanobody** | **Antigen** | | |
| 6LR7 | 4OBE | NLKSTKGDLQF | Chain A: (26, 'N'), (27, 'H'), (28, 'F'), (29, 'V'), (30, 'D'), (31, 'E'), (32, 'Y'), (33, 'D'), (34, 'P'), (35, 'T'), (36, 'I'), (37, 'E'), (38, 'D'), (39, 'S'), (40, 'Y'), (41, 'R') |
| | | TTGNGKRYR | Chain A: (58, 'T'), (59, 'A'), (60, 'G'), (61, 'Q'), (62, 'E'), (63, 'E'),(64, 'Y'), (65, 'S'), (66, 'A'), (67, 'M'), (68, 'R'), (69, 'D'), (70, 'Q'), (71, 'Y'), (72, 'M'), (73, 'R'), (74, 'T'), (75, 'G'), (76, 'E') |
| 6LR7 | 5LRT | NHFVDEYDPTIEDSYR | Chain A: (38, 'N'), (39, 'L'), (40, 'K'), (41, 'S'), (42, 'T'), (43, 'K'), (44, 'G'), (45, 'D'), (46, 'L'), (47, 'Q'), (48, 'F') |
| | | TAGQEEYSAMRDQYMRTGE | Chain A: (160, 'T'), (161, 'T'), (162, 'G'), (163, 'N'), (164, 'G'), (165, 'K'), (166, 'R'), (167, 'Y'), (168, 'R') |
| 7EOW | 8PWH | DQCVACAHYKDPPFCVARCP-SGVKPDLSYMPIWKFPDEEGACQP | Chain E: (583, 'K'), (555, 'F'), (603, 'P'), (579, 'P'), (592, 'W'), (569, 'K'), (572, 'P'), (582, 'V'), (570, 'D'), (560, 'D'), (557, 'P'), (571, 'P'), (561, 'Q'), (585, 'D'), (558, 'E'), (602, 'Q') |
| | | LHCPALVTYNTDTFESMPNP-EGRYTFGASCVTACPYNYLSTDV | Chain E: (295, 'L'), (329, 'R'), (252, 'Y'), (255, 'D'), (257, 'F'), (284, 'T'), (311, 'K'), (285, 'D'), (235, 'H'), (128, 'K'), (248, 'A'), (294, 'P'), (297, 'N'), (268, 'T'), (296, 'H'), (290, 'T'), (286, 'V'), (236, 'F'), (245, 'H'), (254, 'T') |

#### S6.1 Sequence diversity of generated nanobodies

To assess the ability of each CDR design method — ADesigner, dyMEAN, and DiffAb — to generate diverse designs across different binding interfaces, we compared the sequence diversity of their CDRH3 nanobody designs. For consistency, all tools used the same set of docked models during inference, i.e., we did not use the NanoDesigner workflow but the methods directly. We generated sequence logos using the Logomaker library [7], where the $x$-axis represents the length of the CDRH3 sequences derived from PDB structures, and the $y$-axis shows the number of models included in the analysis. For illustrative purposes, we show a small subset of the generated CDR loops.

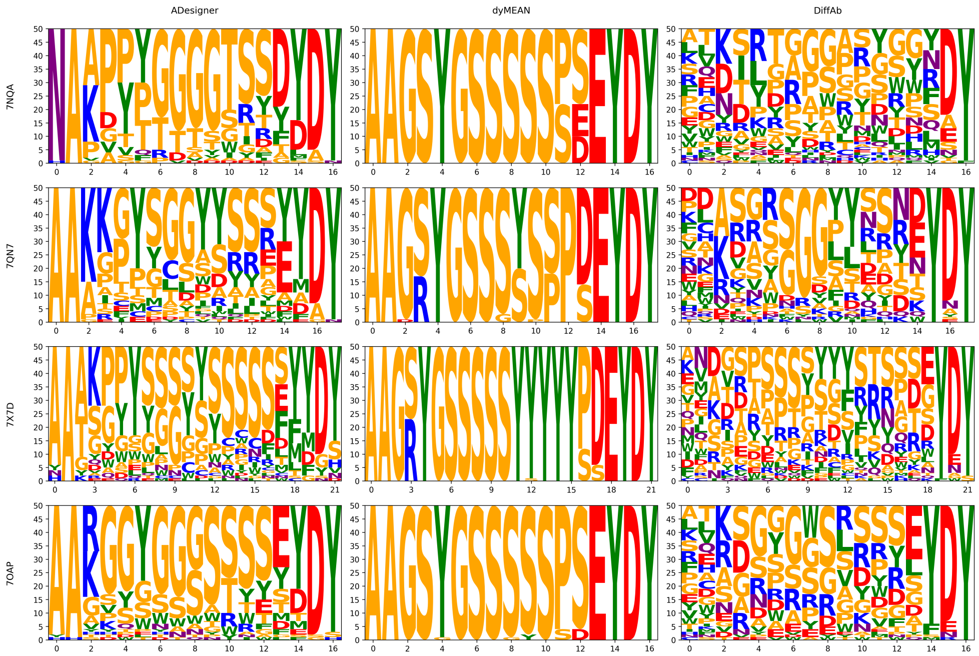


**Fig. S6**: CDRH3 sequence diversity of designed CDRH3 loops across methods for four cases. For entry `PDB:7OAP`, the nanobody referred to as "C1 nanobody" in the Protein Data Bank was used as input for the NanoDesigner process.

As shown in Figure S6, dyMEAN produced nearly identical sequences, indicating potential overfitting and limited generative capacity. ADesigner generated more diverse sequences but tended to include repetitive amino acids. DiffAb exhibited the highest sequence variability, indicating a broader exploration of the design space. This suggests a potential for generating diverse interactions.
